# A qPCR assay for the detection of *Phytophthora abietivora*, an emerging pathogen on fir species cultivated as Christmas trees

**DOI:** 10.1101/2024.09.03.610995

**Authors:** Guillaume Charron, Marie-Krystel Gauthier, Hervé van der Heyden, Guillaume J. Bilodeau, Philippe Tanguay

## Abstract

Emerging species of the *Phytophthora* genus are among the most important threats to global plant biodiversity. For instance, *Phytophthora* root rot (PRR) of Christmas trees is responsible for 10% of the observed mortality rate in nurseries. Diagnosis of PRR involves isolation followed by morphological and molecular identification of the causal agents. However, these methods are rarely adapted to larger scale experiments such as *in situ* detection. For these applications, molecular detection of environmental DNA (eDNA) provides the high-throughput and the fast result generation needed. *Phytophthora abietivora* was associated to PRR in firs cultivated as Christmas trees in the province of Québec (Canada). This study focused on developing a sensitive and specific qPCR assay targeting *P. abietivora* and validating its efficiency on eDNA samples. A set of primers and probe was designed for this assay, and parameters such as the limit of detection (LoD_95%_) and limit of quantification (LoQ) were measured. The assay was tested on eDNA obtained from healthy-looking and PRR symptomatic firs. The assay was shown to be semi-specific because it cross-reacted with P. abietivora, and four phylogenetically close species unrelated to fir diseases. The limit of detection (LOD_95%_) was estimated at 10 copies per reaction (C_q_ of 35.7). The assay showed reliable detection down to 33 *P. abietivora* oospores per gram of soil. Out of 488 eDNA samples from soil, 68 tested positive for *P. abietivora.* While factors such as the tree species, the sampled region or the year of sampling did not affect the proportion of positive results, samples from trees showing PRR-like symptoms had significantly higher odds of testing positive compared to healthy-looking trees. This assay will be useful for rapid diagnostics of *P. abietivora* infected trees and as a prospecting tool to better characterize the natural distribution and dissemination of the disease.

## Introduction

Emerging plant pathogens have always threatened worldwide agriculture, natural ecosystems, and downstream human health [1,2]. Of these, species of the *Phytophthora* genus, with over 200 named species, represent the most destructive plant pathogens known, responsible for historical economic losses [3–7]. *Phytophthora* root rot (PRR) of Christmas trees is one of the most prevalent and widely distributed diseases of true firs, killing about 10% of the planted Christmas trees in the United States of America [8]. Disease symptoms such as root rot, wilting, and chlorotic foliage are hardly distinguishable from those caused by abiotic stresses and other soilborne pathogens such as *Pythium* and *Fusarium spp*. Current diagnosis of PRR in firs requires laborious isolation and identification of the causal agent, which is further challenged given the secondary colonization of infected root and the outcompeting growth of other fungi and oomycetes on semi-selective isolation media [7]. Direct molecular detection of *Phytophthora spp*. from diseased plant material and soil samples has relied lately on conventional PCR amplification of the internal transcribed spacer (ITS) of the ribosomal DNA (rDNA), nested PCR, followed by Sanger sequencing, and qPCR [9–12]. While detection is usually specific, the identification down to the species level can be hampered by the concomitant presence of multiple closely related *Phytophthora spp*. in the samples under investigation [13].

While *P. cinnamomi* is the main species causing PRR in the eastern United States [14], a recent survey identified *P. abietivora* as the species most frequently associated with dying Balsam and Fraser firs cultivated as Christmas trees in Québec, Canada [15]. *Phytophthora abietivora* is a recently described oomycete pathogen isolated in a Connecticut fir plantation and confirmed as a causal agent of PRR in Fraser fir [16]. This species has also been found in the Pennsylvania Department of Agriculture’s historical collection [14]. Because *P. abietivora* have been identified from samples pre-dating its formal description, Molnar et al. [14] suggested that due to the ITS sequence being highly conserved between *P. abietivora* and *P. europaea*, *P. abietivora* isolates could have been misidentified as *P. europaea* for decades. This high sequence identity for the ITS was also found between *P. abietivora* and four other species from clade 7a: *P. uliginosa*, *P. flexuosa*, *P. tyrrhenica* and *P*. sp. *cadmea* [17]. Bily et al. [11] reported associations between *P. abietivora* and tissues symptomatic for *Phytophthora* infections from multiple deciduous tree species. Based on these observations, *P. abietivora* could have been in North America for decades and is now emerging as a pathogen with the potential to cause diseases in both deciduous and evergreen trees. Therefore, developing a molecular assay that detects *P. abietivora* in environmental or diseased tissues is of utmost importance for both biomonitoring and diagnostic applications.

Specific identification and quantification of plant pathogens are now commonly achieved with reliable qPCR assays [7,18]. The high sensitivity makes it a valuable tool for early detection of pathogens, even before disease symptoms are visible [19]. As such, many studies have developed specific qPCR assays targeting *Phytophthora* agents, such as *P. cryptogea* [20], *P. ramorum* [9], *P. cactorum* [21] and many others [22,23]. The need to develop an early and accurate molecular detection assay to monitor *P. abietivora* has become crucial, as treatment options are limited, and prevention remains the best option for growers.

The objectives of this study were to develop a sensitive and specific qPCR assay targeting *P. abietivora,* and to validate its efficiency on infected plant material and soil samples collected during our 2019-2021 survey of PRR in Christmas tree plantations in Québec, Canada.

## Material and Method

### Quantitative PCR primers and probe design

The internal transcribed spacers (ITS) ITS1 and ITS2, including the 5.8S ribosomal RNA region of the rRNA cistron, was chosen as the target gene for our detection assay, as it is commonly used as a barcode for *Phytophthora* species. To design qPCR primers specific to *P. abietivora,* 14 sequences from various *Phytophthora* species isolated from Christmas tree plantations, including *P. abietivora*, were gathered from the GenBank database [15,24]. A sequence from an undescribed *Phytophthora* isolate obtained in a previous study [15] was also included. Fourteen sequences from other *Phytophthora* never reported in association with fir trees were also added, three of which (*P. uliginosa*, *P. flexuosa* and *P. tyrrhenica*) are part of an unresolvable genetic cluster for the ITS1 sequence along with *P. abietivora* and *P. europaea*. The sequences were aligned using Clustal X [25], and the alignment was reviewed for variations to identify potential target regions. Initial specificity assessment was made by querying the GenBank database using the BLAST Sequence Analysis Tool [26] with the sequences of the selected primers and probe (S1 Table).

### Real-time qPCR conditions

A QuantStudio 5 instrument (Thermo Fisher Scientific, Waltham, United States) was used to determine the specificity and sensitivity of the qPCR assay. The detection from environmental samples was done with both a QuantStudio 5 and a 7500 Fast qPCR (Thermo Fisher Scientific, Waltham, United States) instruments. Primers and probe sequences are listed in Table 1 and were synthesized by Integrated DNA Technologies Inc. (Coralville, IA, USA). Cycling conditions were as follows: 5 minutes at 95°C followed by 40 cycles of 15 seconds at 95°C and 45 seconds at 60°C. A single reaction mix comprises 1× SensiFAST™ Probe No-ROX mix (FroggaBio, Concord, Canada), 500 nM of P_abi_ITS_F and P_abi_ITS_R primers, and 100 nM of P_abi_ITS_probe. Unless stated otherwise, the total reaction volume for a standard P_abi_detect detection assay is 10 µL.

**Table 1.**
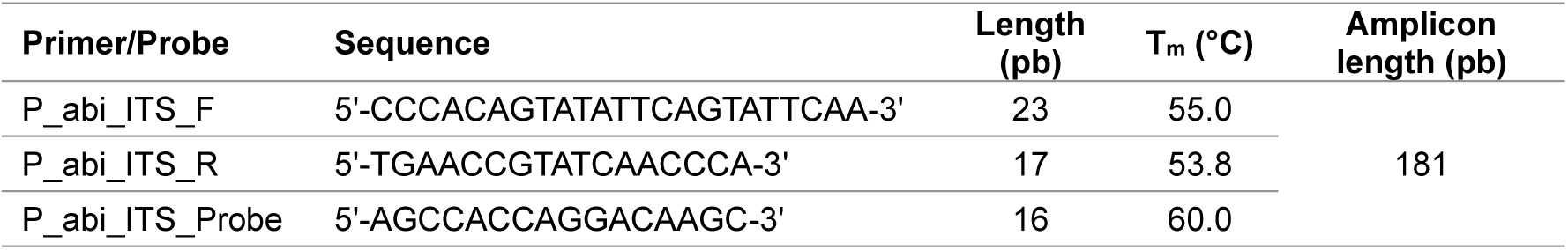
Primers and probe used in the *P. abietivora* detection assay.

### Assay specificity

The specificity of the assay was tested with the genomic DNA from 24 of the 29 species used in the alignment for primer design (no DNA was available for *P. flexuosa, P. uliginosa, P. tyrrhenica, P. pseudotsugae* and *P. citrophthora*). The DNA was obtained from existing in-house collections or was shared by collaborators (S2 Table). Some DNA samples had known DNA concentrations either quantified with a NanoDrop ND-1000 Spectrophotometer instrument (Thermo Fisher Scientific, Waltham, United States) or a Qubit dsDNA assay kit (Thermo Fisher Scientific, Waltham, United States). These samples were diluted to 100 pg/µL for the specificity assay. Other samples had a known concentration in copies (5,000) of Beta-tubulin determined by SYBRGreen real-time PCR quantification [27]. These samples were diluted to 500 copies/µL. For each sample, the assay was run in triplicate on a single 96-well plate. DNA extracted from a purified strain of *P. abietivora* was used as a positive control and PCR-grade water was used as a negative control [15]. To be considered positive, a qPCR reaction had to generate an amplification curve with a C_q_ lower than the cut-off values determined with the LOD_95%_ in all three replicates, as detailed in the results section.

In addition to the panel of species, a panel of 10 *P. abietivora* isolates and 21 *P. europaea* isolates was also tested (S3 Table). The DNA from *P. abietivora* isolates was extracted from purified mycelium, while the DNA from *P. europaea* isolates was extracted from a small quantity of mycelium on agar plugs. The DNA extracts concentrations were measured with a NanoDrop ND-1000 Spectrophotometer (Thermo Fisher Scientific, Waltham, United States). The extracts were diluted to 100 pg/µL, and 1 µL of was used in three technical replicates of the assay. For isolates with failed amplification at 100 pg/µL, the assay was rerun with 1 µL at 500 pg/µL.

### Limit of detection (LOD)

To assess the 95% limit of detection (LOD_95%_) of the assay, a gBlock™ (Integrated DNA Technology Inc., Coralville, IA, USA), including the 181 bp of our target sequence with 24 bp flanking regions on each side, for a total of 229 bp, was designed (S4 Table). The copy number of the gBlock™ stock solution was estimated using the following equation:

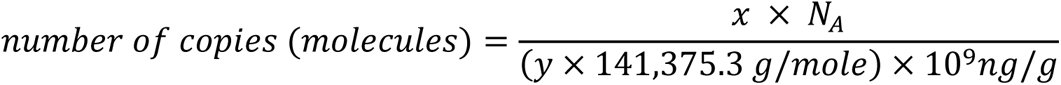

Where x is the amount of the *P. abietivora* ITS2 gBlock™ in nanograms, N_A_ is the Avogadro constant (6.0221×10^23^), and y is the length of the *P. abietivora* ITS2 gBlock™ in bp.

A solution at 10 million copies per microliter was prepared and diluted tenfold seven times to obtain 10^6^ to 1 copy per microliter dilutions. One microliter of each dilution was used in three technical replicates of the assay. Because the 1 and 10 copies dilutions had great variations in their C_q_ values, only the dilutions from 100 copies to 10^6^ copies were used to calculate the standard curve parameters. The standard curve equation was determined by linear regression analysis using the Log_10_ number of copies as the predictor variable, and the C_q_ value as the response variable. The efficiency of the amplification was calculated as:

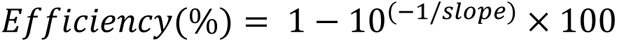

Once the standard curve was established, the LOD_95%_ was tested with twofold serial dilutions of the gBlock™, including 20, 10, 5, 2.5, 1.25, and 0.625 copies per microliter. For each dilution, 5µL was used in 20 technical replicates of the assay (100, 50, 25, 12.5 6.25 and 3.1 copies per reaction). A successful amplification was defined as any amplification with a C_q_ value under 40. The theoretical LOD_95%_ was determined with a binary logistic regression (probit model) analysis using the Log_10_ of the number of copies as the predictor variable, and the positive/negative detection result as the response variable. The minimal number of copies necessary to obtain a positive result in 95% of the assays (LOD_95%_) was determined with the resulting binary logistic regression equation. The LOD_95%_ value was empirically verified by making a dilution of the gBlock™ to the determined LOD_95%_ value and tested in 20 replicates.

### Limit of quantification (LOQ)

To estimate the limit for *P. abietivora* quantification in environmental samples, oospore suspensions of different concentrations were prepared. The *P. abietivora* isolate CFL062-E was inoculated on a V8-Phytagel medium (5% V8 juice, 0.4% Phytagel^TM^ (MilliporeSigma, Burlington, USA) amended with 10% of a macro-element solution (10 g/L KH_2_PO_4_, 5 g/L MGSO_4_•7H_2_O and 1 g/L CaCl_2_•2H_2_O) and incubated at 20°C. After 7 days, the mycelium obtained was used to inoculate five new V8-Phytagel^TM^ media that were incubated for another 7 days at 20°C. Following incubation, the Phytagel^TM^ media were dissolved in 1L of citrate buffer (10 mM, pH 6) and filtered on 100 µM nylon mesh to gather the *P. abietivora* mycelium. To free the oospores, the collected mycelium was resuspended in 30 mL of an enzyme solution containing 10 mg/mL Driselase™ Basidiomycetes sp. (MilliporeSigma, Burlington, USA) and 6.7 mg/mL of lysing enzymes from *Trichoderma harzianum* (Millipore Sigma, Burlington, USA) and incubated overnight at 30°C on a tube rotator (VWR International, Radnor, USA). Complete digestion of the mycelium was confirmed by microscopic observation. The oospore suspension was centrifuged 5 minutes at 2800 g, and the supernatant was removed. The oospore pellet was washed twice in phosphate-buffered saline (PBS, pH 7.4), and resuspended in 1 mL of PBS. Oospore count was estimated using an Improved Neubauer hemocytometer.

For DNA extraction without soil, the oospores were resuspended in 180 µL of ATL buffer from the QIAamp DNA mini kit (Qiagen, Valencia, CA, USA). For DNA extractions with soil, the oospores were added to a PowerBead pro tube containing 250 mg of garden soil exempt from *P. abietivora*. In both cases, DNA extractions followed the protocol provided by the manufacturer and elution was performed in 100 µL of PCR grade water. Following extraction, 1 µL of each DNA sample was used with the *P. abietivora* detection assay in three technical replicates, providing data for DNA equivalent of 2 to about 33,000 oospores per gram of soil. After data compilation, the Hampel filter method was used to identify and remove potential outlier values from the final data set [28]. This filter considers as outliers the values outside the interval formed by the median, plus or minus 3 median absolute deviations. The limit of quantification was established as the lowest number of oospores, with a 100% positive detection rate over the replicates left after outlier removal.

### Environmental samples preparation

Soil associated with trees of three different statuses (PRR-like symptomatic trees from Christmas tree plantation, asymptomatic trees from the same plantation, and trees from natural forest nearby the plantation) were sampled in a previous study [29] from 40 sites across the Estrie and Chaudière-Appalaches regions (Québec, Canada) in between 2019 and 2021. Unidentified root fragments were sampled only from the trees showing PRR-like symptoms. The soil and root samples were prepared as described in Charron et al [15]. In summary, the soil samples were mixed by hand and sieved through a 4-mm mesh basket. From this homogenized soil, approximately 250 mg were transferred to a 2 mL tube for DNA extractions. Root fragments retained in the sieve were cut into 5 to 10-mm sections with a sterile scalpel and transferred to a 2 mL tube for DNA extractions.

### DNA extractions

DNA from soil and root fragments (Estrie and Chaudière-Appalaches) was extracted with the DNeasy® Powersoil kit (QIAGEN, Hilden, Germany, discontinued) with slight modifications. Instead of 10 minutes, the samples were vortexed to maximum speed for 20 minutes. PCR grade water was used for the elution of DNA instead of the C6 Buffer. DNA from purified mycelium from *Phytophthora* isolates and from *Phytophthora* mycelium on agar plugs was extracted using the protocol developed in Penouilh-Suzette et al. [30] with minor modifications as in Charron et al. [15]. DNA for each DNA sample obtained from soil, roots, and baiting leaves, 1 µL was tested for the presence of *P. abietivora* with the P_abi_detect assay in three technical replicates.

### Data visualization and Statistical analyses

All data visualization was made with the help of the R software version 4.1.3 [31] and custom scripts. Unless stated otherwise, the significance level used for all statistical tests is *ɑ*= 0.01 and p-values were adjusted with the false discovery rate method when appropriate (fdr) [32]. Maps were generated with the package sf v1.0-16 [33] and SHP files publicly available from the Geographic and administrative database [34]. Linear and binary logistic regressions were respectively performed with the lm and glm functions of the R software [31].

## Results

### Quantitative PCR design

The target region of our P_abi_detect assay can be found in the ITS2 sequence between position 728 and 917 of our alignment (S1 Table). While the sequence of the primers P_abi_ITS_F and P_abi_ITS_R does not prevent the amplification of nonspecific targets, the assay’s specificity comes from the probe P_abi_ITS_Probe (S1 Table and Table 1). However, the two sister species *P. europaea* and *P. abietivora,* as well as *P. flexuosa, P. tyrrhenica,* and *P. uliginosa* all have 100% sequence identity in the target region. The next closest species to *P. abietivora* in our alignment, *P. fragariae*, has two SNPs in the sequence targeted by the probe (S1 Table).

### Assay specificity

Criteria for positivity were first established as follows: to be considered positive, a sample had to produce an amplification curve for all three replicates, with a C_q_ lower than 35.7, a value determined by the experimental LOD_95%_ (see section on LOD). As expected, the P_abi_detect qPCR showed specific amplification when 100 pg genomic DNA from *P*. *abietivora* or *P*. *europaea* was used in tests (Fig 1A). No nonspecific amplification under the LOD_95%_ cut-off was observed in the tested conditions for the genomic DNA of the 22 other species tested, including members of subclade 7 *P*. *fragariae*, *P.* ×*cambivora*, and *P. cinnamomi*. Following the criteria for positivity established above, three species (*P*. *cactorum*, *P*. *sansomeana* and the unknown *Phytophthora*) amplifying at a late cycle, past the cut-off and only for one replicate out of three, were not considered as positive and may be the result of a cross-contamination.

**Figure 1.**
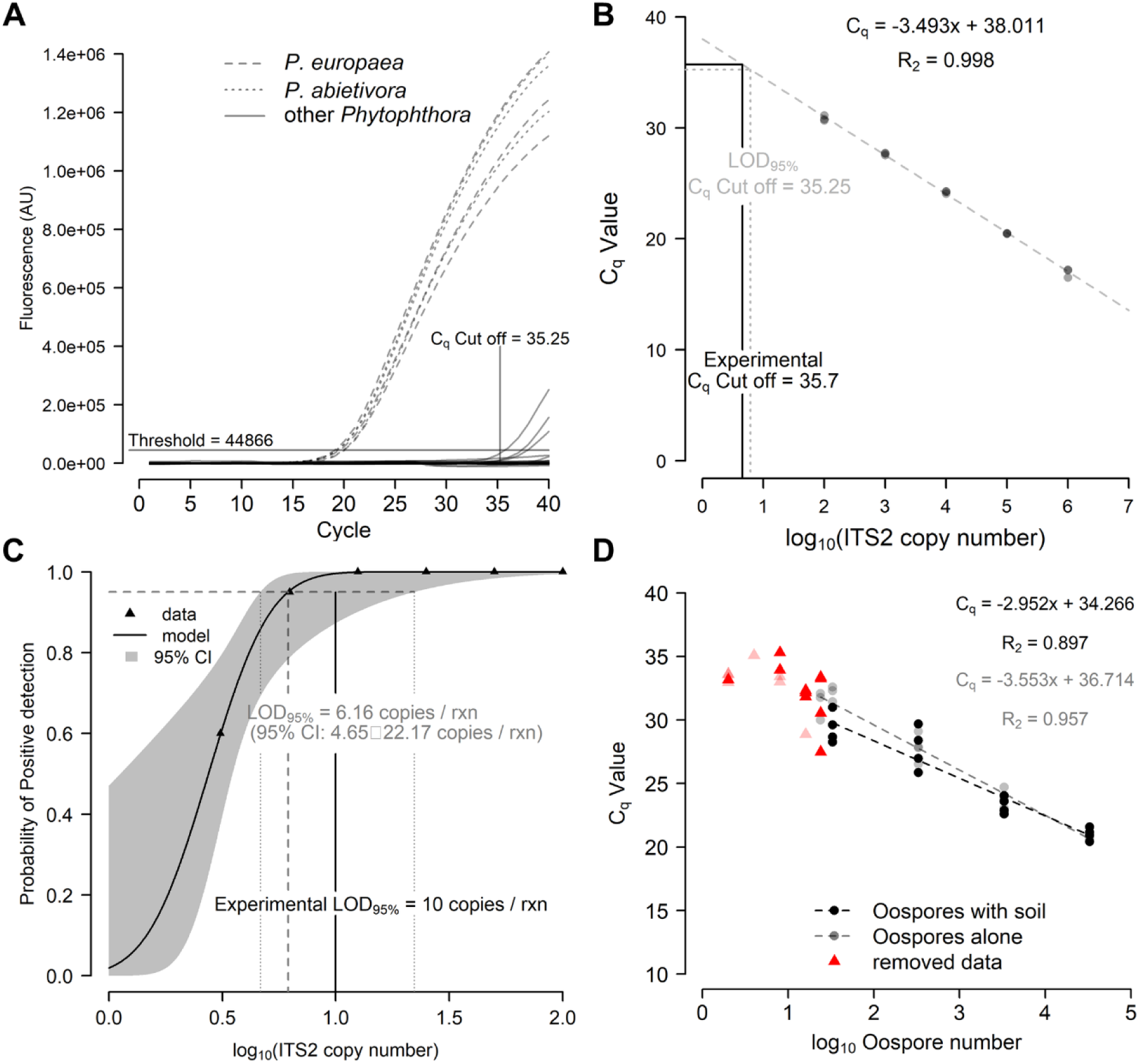
The P_abi_detect assay is specific and sensitive. (A) The P_abi_detect qPCR assay shows specific amplification from *P. abietivora* and *P. europaea* genomic DNA while other species DNA do not result in reliable amplification under the established cut-off. (B) The P_abi_detect assay shows a linear response between C_q_ values and the ITS2 copy numbers. (C) The Probit regression analysis used to estimate the LOD95% of the assay. The dashed lines intercept at the theoretical 95%. (D) The P_abi_detect shows a linear response between C_q_ values and the quantity of oospores present in samples spiked with known quantities of cells.

Out of 31 independent *P. abietivora* and *P. europaea* isolates DNA tested, 29 (nine *P. abietivora* and 20 *P. europaea*) consistently tested positive with an average C_q_ value of 23.1 (S3 Table). The variation in C_q_ between samples was high, which could be due to the initial dosage of DNA. Two DNA samples, one *P. abietivora* and one *P. europaea*, produced less than three positive replicates and were retested to verify if this was due to an overestimation of the DNA concentration due to poor quality samples. The samples were retested at 500 pg of DNA per reaction. The assay gave a clear positive signal for the *P. europaea* DNA samples (AX44). The *P. abietivora* sample (12324_2CR) also failed to produce a positive detection at 500 pg. The identity of the species was verified by Sanger sequencing of the whole ITS1-5.8S-ITS2 region. A BLAST of the obtained sequence against the nt/nr database of the NCBI confirmed a misidentification as the DNA did not match to *P. abietivora*, but *P. gonapodyides*.

### Standard curve

Our assay has shown a linear response between 1 million target gBlock™ copies down to 100 target gBlock™ copies per reaction (Fig 1B). The amplification efficiency of 93.3% suggests that PCR products are not exactly doubling at each cycle. However, this value is within the desired efficiency range which is between 90 and 110% [35,36].

### Limit of detection (LOD)

We evaluated the analytical sensitivity of the P_abi_detect qPCR with two-fold dilutions of a positive control gBlock™ between 100 and 3.1 copies per reaction. The binary logistic regression analysis estimated the LOD_95%_ at 6.16 copies per reaction (Fig 1C). The 95% confidence interval was from 4.65 to 22.17 copies per reaction. Repeating the P_abi_detect qPCR assay at 6.16 copies per reaction in 20 replicates yielded only a 75% positive rate. To provide a more reliable analytical cut-off for assay positivity, we tested 8,10, and 25 copies per reaction of the control gBlock™. At 10 copies per reaction, the assay returned a 95% positive rate with an average C_q_ of 35.7 (S5 Table). Extrapolating the C_q_ value from the copy number with the regression line of our standard curve would result in an analytical cut-off of 34.5. However, since these values fall outside the linear part of the standard curve, that 10 copies represent a quantity within the 95% confidence interval, and that false negatives could be more detrimental than false positives, we decided to use the mean C_q_ value obtained for 10 copies per reaction as experimentally tested. This resulted in a more conservative analytical LOD_95%_ of 35.7. In all subsequent experiments, C_q_ values exceeding this cut-off would be considered negative.

### Limit of quantification (LOQ)

We estimated the limit of quantification by adding different suspensions of known oospore quantities, ranging from .5 to about 33,000 oospores, in DNA extractions with soil and without soil. After outlier removal, multiple linear regression was used to test if the quantity of oospores present in a sample significantly predicted the assay C_q_ values for DNA extracted from soil spiked with oospores or oospores alone (Fig 1D). The overall regression was statistically significant (R^2^= 0.937 (multiple) and .931 (adjusted), F(3,30)= 148.3, p< 0.001). Both the Log_10_ of the oospore quantity in soil (β= −2.952, p< 0.001) and pure oospores (β= −3.553, p< 0.001) DNA extracts significantly predicted the C_q_ values. While there was a significant difference between the two intercepts (2.448, p= 0.021), the difference in slope was not significant (-.601, p= 0.07). The lowest number of oospores detected in 100% of the tests was 8.3 or about 33 oospores per gram of soil. The average C_q_ values for this oospore quantity were 29.4 (with soil) and 32.1 (only oospores).

### Case study from plantations

The P_abi_detect assay was used to detect *P. abietivora* from DNA extracted from soil samples taken under 1) healthy-looking firs in natural forest, 2) healthy-looking firs in plantation, and 3) PRR symptomatic firs in plantation (Fig 2). Unidentified root samples were only taken from the rhizosphere soil of PRR symptomatic firs. Of 488 soil samples, 68 were declared positive for *P. abietivora.* No statistically significant difference in the proportion of positive detection was found between *Abies* species sampled (2-sample test for equality of proportion, *p*= 0.013), region sampled (2-sample test for equality of proportion, *p*= 0.069) or years of sampling [Pairwise comparison of proportions, *p*= 0.065 (2020-2021 and 2019-2020) and *p*= 0.953 (2019-2021)]. However, we found a significant difference in the proportion of positive detection between soil sampled from PRR-like symptomatic trees and healthy trees [Pairwise comparison of proportions, p= 5.1×10^-6^ (PRR-healthy forest), p= 0.0008 (PRR-healthy plantation), and p= 0.172 (healthy forest-healthy plantation)] (Fig 3A). Accordingly, a binary logistic regression using the result of the assay (positive, negative) as the dependent variable found that soil eDNA samples from PRR symptomatic trees in Christmas tree plantations had higher odds of being positive for *P. abietivora* (OR = 5.96; 95% CI: 2.77 to 12.85; p = 5.19×10^-6^, Fig 3B) compared to healthy looking trees from natural forests. The odds of obtaining a positive detection result from healthy-looking trees in Christmas tree plantations were also higher than healthy-looking trees from natural forests, but this difference was not statistically significant (OR = 1.92; 95% CI: 0.82 to 4.48; p = 0.13, Fig 3B). For PRR symptomatic trees, the proportion of positive detection did not significantly differ between soil (42/164) or root (28/162) samples (2-sample test for equality of proportions with continuity correction, X^2^(1) = 1.79, p=0.090). There were only 16 samples overlapping for positive detection between both sample types (Fig 4). The average C_q_ values obtained for soil and root samples were also not significantly different (Welch two sample t-test, t(50.19)=1.4911, p=0.14).

**Figure 2.**
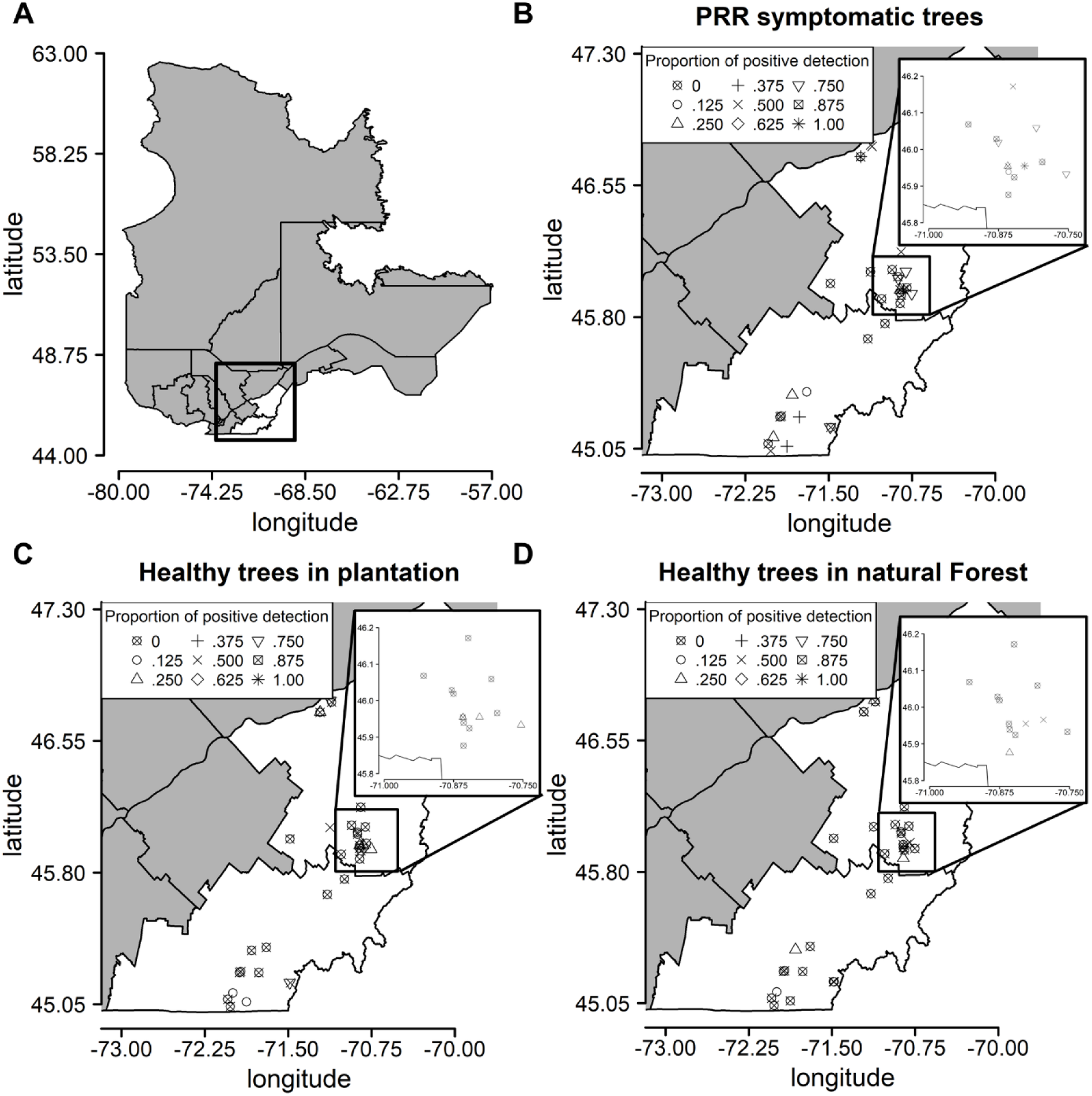
*Phytophthora abietivora* is present in major Christmas tree producing regions of the province of Québec. (A) Map of the province of Québec with the regions of Estrie (Southward) and Chaudière-Appalaches (Northward) highlighted in white. (B) Proportion of positive detections in soil samples of PRR symptomatic firs trees for each sampled site. (C) Proportion of positive detections in soil samples of healthy-looking firs trees for each sampled site. (D) Proportion of positive detections in soil samples of healthy-looking firs trees in natural forest stands in the vicinity of each sampled site. The insets in (B), (C), and (D) represent an enlarged region with a high density of sampled Christmas tree productions.

**Figure 3.**
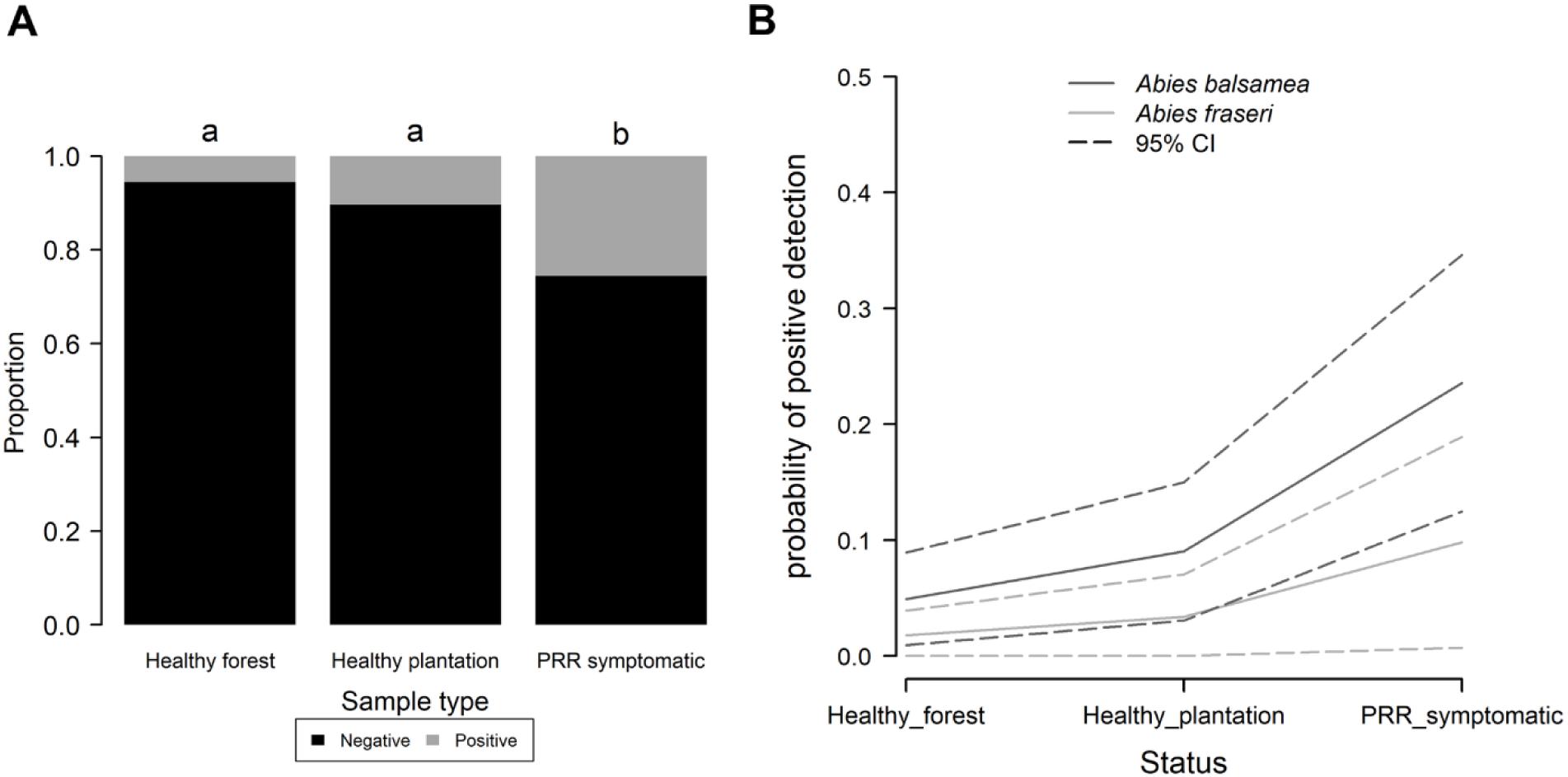
PRR-like symptoms are a predictor of the presence of *P. abietivora* in the associated rhizosphere soil. (A) Bar plot showing the proportion of positive detections for the soil samples of different tree status. The number of samples in each category is indicated above the bars. (B) Binary logistic regression using the result of the assay (positive, negative) as the dependent variable. For both cultivated *Abies* species, samples taken in the rhizosphere soil of PRR-like symptomatic trees have higher odds of being positive for the presence of P. abietivora than the other tree status.

**Figure 4.**
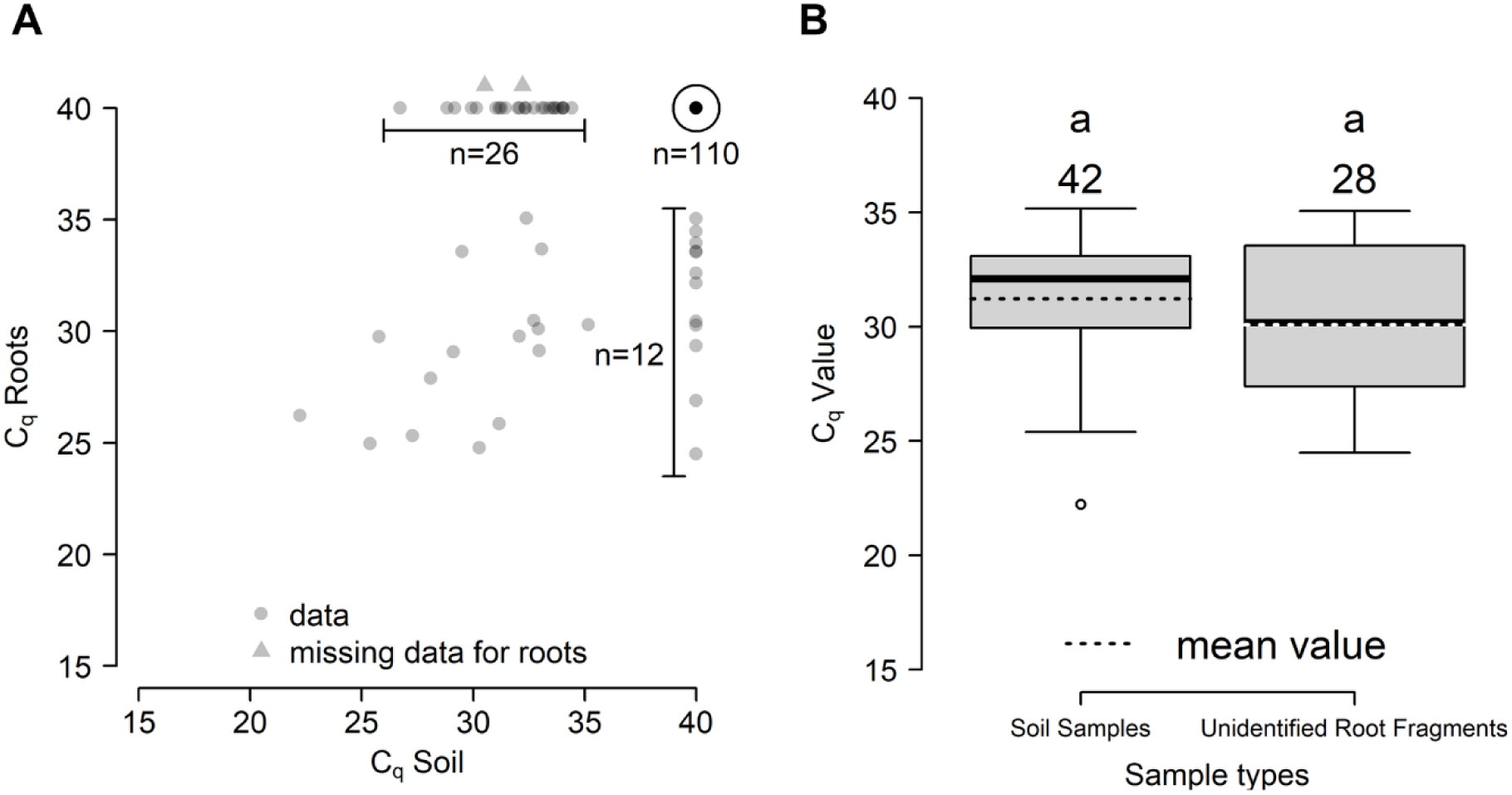
eDNA from *P. abietivora* is found in soil and root fragments samples in comparable quantities. (A) Scatter plot of C_q_ values for the 164 samples from PRR-like symptomatic trees. (B) Boxplot of the C_q_ values obtained for positive soil and unidentified root fragments samples. Number of samples is displayed above the individual plots.

## Discussion

Since *P. abietivora* is a recently described species [16], most of the information about its distribution and host range remains to be discovered. To facilitate and speed up filling these gaps in knowledge, we developed a sensitive and specific qPCR assay to detect *P. abietivora* in environmental samples (diseased plant tissues and soil).

Although the P_abi_detect assay was not designed with quantification in mind, the parameters obtained with the standard curve indicate that the assay can be used in a quantitative manner. Based on the data obtained for gBlock™ of the target sequence, as little as 10 copies of the target sequence should be detectable. While this information was not obtained for *P. abietivora* genomic DNA, the results obtained from the assay, when used on the DNA of different strains, suggest that it is as sensitive as other TaqMan assays developed in the ITS for other *Phytophthora* species [7]. Our assay takes advantage of the high copy number of the rRNA in the *Phytophthora* genomes [37]. This increases the sensitivity of our assay in samples with low pathogen concentration compared to other assays based on the amplification of single-target genes. Even in the case of mature oospores, which contain only one diploid nucleus [38], our assay provided detection at concentrations as low as 33 oospores per gram of soil. Previous studies have shown that the detection of *Phytophthora spp* eDNA from soil is feasible even at those low densities [39,40], while other found it harder to effectively detect Phytophthora pathogens due to DNA extraction limitations [41].

Latent structures such as oospores from *P. capsici* were still viable after two years in soil exposed to natural environmental conditions [42], and for at least five years for *P. abietivora* [16]. This means that our assay could be used to estimate the reduction of *P. abietivora* propagules in soil following disinfection treatments.

In the past, others have targeted different genes to differentiate *Phytophthora* species [43], such as Ypt1 [44], Atp9-Nad9 [22,23] or Rps10 for oomycetes in general [45]. While using the ITS as a target for detection offers advantages towards sensitivity, it does come with a trade-off in specificity. The ITS as a marker provides resolution between distant species, but its use is sometimes challenged when dealing with closely related species [46]. For the P_abi_detect assay, this translates as being only semi-specific as the ITS2 sequence targeted by the primers and probe is identical between the five members of the *P. europaea* species complex, which includes *P. abietivora*, *P. flexuosa*, *P. uliginosa* and *P. tyrrhenica* [17]. This shortcoming of the assay is mitigated by the fact that, besides *P. europaea*, none of the other members of this complex has ever been reported in association with evergreen trees or reported in North America (*P. flexuosa*) [47], (*P. uliginosa*) [48], (*P. tyrrhenica*) [49]. Previous reports of *P. europaea* on North American firs are questionable since there are chances that they resulted from misidentification due to the high identity between the ITS sequences of *P. europaea* and *P. abietivora*. In cases for which isolates are available, sequencing of loci offering more discrimination will be needed for confirmation. The P_abi_detect assay has been developed to provide a tool for detection and diagnosis in North American Christmas tree productions. Remaining within this context excludes the presence of the other species from the *P. europaea* species complex. When the presence of other species from this complex cannot be excluded, this assay should always be coupled with confirmation methods such as strain baiting or strain isolation from diseased tissues, followed by Sanger sequencing of a discriminating loci to confirm the presence of only *P. abietivora*. There is currently limited genetic information for *P. abietivora*, with only 6 different loci sequences available from the GenBank® database. While these sequences offer enough resolution for taxonomic studies, developing specific detection assays will require more sequence information. Whole genome sequencing of *P. abietivora* and the other species from the complex will undoubtedly facilitate the development of more specific detection tools.

We used the P_abi_detect assay on DNA extracted from different samples gathered in Christmas tree plantations during a previous study [15]. In this study, *P. abietivora* was identified as a potential pathogen, with eight strains being isolated from independent PRR-like symptomatic tree samples. Based on isolation data, *P. abietivora* was detected in 4.3% (7 out of 164) of the rhizosphere soil samples and 0.6% (1 out of 164) of the root samples [15]. In contrast, with the P_abi_detect assay, positive detections for rhizosphere soil samples and roots samples increased 5.95-fold (to 25.6%) and 28-fold (to 17%), respectively. These results further reinforce the hypothesis of a causal link between *P. abietivora* and the PRR diseases observed in the Québec Christmas tree plantations. Moreover, we detected *P. abietivora* in the rhizosphere soil of healthy-looking trees from both the plantations (10.4% of samples) and nearby natural forests (5.6% of samples), whereas none of those samples yielded any *P. abietivora* isolates.

A recent study by Van der Heyden et al [17], built on a subset of the samples analyzed in the present study, used a metagenomic approach and showed the presence of the *P. europaea* complex (most probably *P. abietivora*) in 29.9% of the soil samples collected under trees showing PRR-like symptoms. They also showed that 15% of the soil samples collected under healthy trees also presented reads from the complex. As expected, there is a strong positive association between the results of the P_abi_detect assay and the presence of reads from the *P. europaea* species complex for a given sample.

These independent methods both confirm that *P. abietivora* is present in natural forest stands neighboring the plantations. Its association with the rhizosphere soil of balsam firs from natural forests suggests that it could either be a native pathogen of this fir species or that it spread from nearby infected Christmas tree plantation. Since little is known about the current distribution of *P. abietivora*, our detection assay can be used in mapping efforts to establish its presence in diverse environments such as nurseries, Christmas tree farms, and natural forests.

Large differences in sensitivity between baiting and eDNA detection of *Phytophthora* species in soil samples are not uncommon. Using a nested PCR, *P. cinnamomi* was detected in 96% of the soil samples taken at the margin of the disease front, while only 2.2% of the same samples yielded isolates from baiting [50]. This is not surprising as successful baiting relies on the formation of sporangia, release of zoospores from the sporangia, and infection of the bait tissue by the zoospores. However, various microorganisms such as bacteria and fungi present in the soil can potentially interfere with the growth, reproduction, and infectivity of *Phytophthora* species by producing compounds that can inhibit the production of zoospores [51]. While more sensitive, using eDNA only indicates the presence of the pathogen without any information about its viability status. A positive eDNA detection does not equate to living cells in the sample as it was shown that DNA can persist through time in the environment following the death of organisms [52,53]. It was estimated that the DNA of *P. cinnamomi* cells can persist for more than a year in soil [54]. In consequence, culturing should be used in conjunction with our assay to confirm the viability of the pathogens detected. As demonstrated for *P. ramorum*, our assay should also be tested on reverse transcribed RNA to assess its utility to detect and quantify viable propagules of *P. abietivora*.[55]

In conclusion, this qPCR assay will prove useful in the early detection and biomonitoring of *P. abietivora* in Christmas trees plantations. Furthermore, it will facilitate the gathering of crucial information on the distribution of this species in nature, leading to a better understanding of the ecology of this emerging pathogen.

## Acknowledgments

We would like to thank Amélie Potvin and Nathan Benoît for their technical assistance and insightful discussions; Christopher Keeling for his comments on the manuscript; Jean-François Légaré, Éric Dussault, Christian Lacroix, Dominique Choquette and Jacinthe Drouin for their contributions to fieldwork and sampling; and all the growers for giving us access to their plantations.

## Supporting information

**S1 Table. Alignment of the targeted sequences for different Phytophthora species.**

**S2 Table. DNA from *Phytophthora* isolates used to assess the specificity of the P_abi_detect assay.**

**S3 Table. DNA from *P. abietivora* and *P. europaea* isolates used to test the P_abi_detect assay.**

**S4 Table. Sequence of the gBlock^TM^ fragment used for the standard curve of the P_abi_detect assay.**

**S5 Table. Data obtained for the LOD_95%_ experimental validation. S1 File. Data and script used for analysis.**

