## Supplementary material for "A qPCR assay for the detection of *Phytophthora abietivora*, an emerging pathogen on fir species cultivated as Christmas trees": S1 File: qPCR_analysis_Pabi_detect_2024_08_23.R

```
##Set working directory
setwd("P:/Bureau_desktop_2024_06_21/Detection")

##load needed libraries
library("sf")


########################################
########Data analysis qPCR assay########
########################################

######Amplification plots for the Specificity test
ampdata=read.csv("amplification_data_specificity.csv", header=T)
marks=seq(0,1400000, 200000)

setwd(paste(strsplit(getwd(),"/Data")[1]))
dir.create(paste(getwd(), "Figures", sep="/"))
setwd(paste(getwd(), "Figures", sep="/"))

png("Figure_1A.png", height=2000, width=2000, res=300)
par(mar=c(4,6,4,2))
plot.new()
plot.window(xlim=c(0,40), ylim=c(0,1400000))
axis(1, lwd=2, cex.axis=1.5, at=seq(0,40,5))
axis(2, at=marks, format(marks, scientific=T), lwd=2, las=2, cex.axis=1.2)
title(xlab="Cycle", cex.lab=1.5, line=2.5)
title(ylab="Fluorescence (AU)", line=5, cex.lab=1.2)

for (i in unique(ampdata$Well)){

	sub=subset(ampdata, ampdata$Well == i)
		if(sub$Sample[1] == "M9554"){
		lines(sub$dRn,lwd=2, col=rgb(0,0,0,0.4), lty=2)
		}else if (sub$Sample[1] == "RC2017_1"){
		lines(sub$dRn,lwd=2, col=rgb(0,0,0,0.4), lty=3)
		}else{
		lines(1:40, sub$dRn,lwd=2, col=rgb(0,0,0,.4))
		}
}
segments(35.25,-1,35.25,400000, col=rgb(0,0,0,0.5), lwd=2)
segments(-1,44866, 40, 44866, col=rgb(0,0,0,0.5), lwd=2)
text(6,80000, "Threshold = 44866", col="black", cex=1.2)
par(xpd=NA)
text(35.25,420000, expression("C"[q]*" Cut off = 35.25"), col="black", cex=1.2)
par(xpd=F)
legend("top", c(expression(italic("P. europaea")),expression(italic("P. abietivora")), expression('other ' *italic('Phytophthora')* '')), text.font=c(3,3,1), lwd=2, bty="n", col=rgb(0,0,0,.5), lty=c(2,3,1), cex=1.5)
dev.off()
setwd(unlist(strsplit(getwd(), "/Figures"))[1])


#####probit analysis for LOD95
setwd(paste(getwd(), "Data", sep="/"))
data_LOD=read.csv("LOD_data.csv")
setwd(unlist(strsplit(getwd(), "/Data"))[1])

data_LOD$Log10_copy=log(data_LOD[,1], 10)
data_LOD$Detection=as.integer(data_LOD[,2]<40&data_LOD[,2] != "Undetermined")

copies=unique(data_LOD[,3])

#make a table representing the positive and negative results (positive determined as ct<40)
pos_result=table(data_LOD[,2]<40&data_LOD[,2] !="Undetermined", round(data_LOD[,3],3))
p.pos_result=prop.table(table(data_LOD[,2]<40&data_LOD[,2] !="Undetermined", round(data_LOD[,3],3)),margin=2)

####probit model####

#generate the model with the data from detection in function of the copynumber of target
prob_LOD=glm(Detection~Log10_copy, family=binomial(link="probit"), data=data_LOD)
data_LOD$Predicted_prob=predict(prob_LOD, type="response")

#generate predictions from the model for Log10_copy values between 0 and 2 with 0.5 copy number increments
Log10_new=data.frame(Log10_copy=seq(0,2,length=2000))
Log10_new$predicted_value=predict(prob_LOD,Log10_new, type="link")

#get the invlink function from the model to help create the 95% confidence interval
fam=family(prob_LOD)
ilink=fam$linkinv
linkf=fam$linkf


#get the fitted and se values on the link scale from the model
LODfitted=unlist(predict(prob_LOD,Log10_new, se.fit=T)[1], use.names=F)
LODse=unlist(predict(prob_LOD,Log10_new, se.fit=T)[2], use.names=F)

##calculate the upper and lower ci
lwr_ci=LODfitted - (2*LODse)
upr_ci=LODfitted + (2*LODse)

##find the Log10 copy number where the probability is 95% of positive detection
p95=linkf(0.95)
p95_log10CN=unname((p95-prob_LOD$coefficients[1])/prob_LOD$coefficients[2])
#0.789289
##return the copy number value
10^p95_log10CN
#6.155863

##find the 95% CI copy number values on the LOD95% values
#find the values of the CI curves which are closest to .95
which(abs(ilink(lwr_ci)-.95000) == min(abs(ilink(lwr_ci)-.95000)))
#1346
cimax=Log10_new$Log10_copy[1346]
#1.345673

which(abs(ilink(upr_ci)-.95000) == min(abs(ilink(upr_ci)-.95000)))
#668
cimin=Log10_new$Log10_copy[668]
#.6673337

#Plot the datapoints, the predicted curve and the 95% confidence interval on prediction

setwd(paste(getwd(), "Figures", sep="/"))
png("Figure_1C.png", height=2000, width=2000, res=300)
par(mar=c(5,5,4,1))
plot.new()
plot.window(xlim=c(0,2), ylim=c(0,1))
title(xlab=expression("log"[10]*"(ITS2 copy number)"),
 ylab="Probability of Positive detection", cex.lab=1.25)
axis(1, cex.axis=1.3, lwd=2)
axis(2, las=2, cex.axis=1.3, lwd=2)
points(rev(copies), p.pos_result[2,], pch=17, col="black")
lines(Log10_new$Log10_copy ,ilink(LODfitted), lwd=2)

segments(c(p95_log10CN,0),c(.7,.95),c(p95_log10CN,cimax),c(.95,.95), lty=2, lwd=2, col="grey50")
segments(c(p95_log10CN,p95_log10CN),c(.55, .15),c(p95_log10CN, p95_log10CN),c(.21, -1), lty=2, lwd=2, col="grey50")
segments(c(cimin, cimax, cimin, cimax),c(.95, .95, .15,.15),c(cimin, cimax, cimin, cimax),c(.7, .7,-1,-1), lty=3, col="grey50")
segments(c(cimin, cimax),c(.55, .55),c(cimin, cimax),c(.21, .21), lty=3, col="grey50")
segments(c(1, 1),c(.95, .95),c(1, 1),c(.7, .7), lty=1, lwd=2, col="black")
segments(c(1, 1,1),c(.55, .55, .15),c(1, 1, 1),c(.21, .21, -1), lty=1, lwd=2, col="black")


polygon(append(Log10_new$Log10_copy, rev(Log10_new$Log10_copy)),
ilink(append(upr_ci,rev(lwr_ci))), col=rgb(0,0,0,0.25), lty=0)

legend(0,.95, c("","","95% CI"), fill=c("white","white",rgb(0,0,0,0.25)), bty="n", border="white", cex=1.2)
legend(0,.95, c("data","model"), pch=c(17,NA),
lty=c(0,1), lwd=2, bty="n", cex=1.2)
text(cimin+.45,.65,expression("LOD"["95%"]*" = 6.16 copies / rxn"), cex=1.3, col="grey50")
text(cimin+.6, .6, "(95% CI: 4.65�22.17 copies / rxn)", cex=1.3, col="grey50")
text(1.2,.175, expression("Experimental LOD"["95%"]*" = 10 copies / rxn"), cex=1.3, col="black")
dev.off()
setwd(unlist(strsplit(getwd(), "/Figures"))[1])


######Standard curve

###read data
setwd(paste(getwd(), "Data", sep="/"))
stdcurve=read.csv("Std_curve_data.csv", header=T)
setwd(unlist(strsplit(getwd(), "/Data"))[1])


###get the datapoints
dp=cbind(stdcurve$Sample[1:15], as.numeric(stdcurve$Cq)[1:15])

####calculate the linear regression for the datapoints and the r2
stcurve=lm(dp[,2]~log10(dp[,1]))
rsq=round(cor(dp[,2],log10(dp[,1]))^2, 3)
slope=round(stcurve$coefficients[2],3)
ordori=round(stcurve$coefficients[1],3)

###calculate the ct corresponding to the LOD95 (.789289)
lod95cq=unname(p95_log10CN*stcurve$coefficients[2]+stcurve$coefficients[1])


####extrapolate the curve from 0 to 7 log10 copies
pts=c(0:7)
cqcurve=pts*stcurve$coefficients[2]+stcurve$coefficients[1]

setwd(paste(getwd(), "Figures", sep="/"))
png("Figure_1B.png", height=2000, width=2000, res=300)
par(mar=c(4,5,4,4))
plot(log10(dp[,1]),dp[,2], xlim=c(0,7) , ylim=c(0,40), axes=F, col=rgb(0,0,0,.35), pch=16, xlab=NA, ylab=NA, cex=1.2)
axis(1, at=c(0:7), lwd=2, cex.axis=1.2)
axis(2, at=seq(0,40,10), lwd=2, cex.axis=1.2, las=2)
title(xlab=expression("log"[10]*"(ITS2 copy number)"), cex.lab=1.5, ylab=expression("C"[q]*" Value"))
lines(x=pts, y=cqcurve, lty=2, lwd=2, col=rgb(0,0,0,0.25))
text(4,36, bquote(paste("R"["2"]*" = ",.(rsq))), cex=1.3) 
text(4,39, bquote(paste("C"["q"]* " = ", .(slope), "x + ",.(ordori))),  cex=1.3)
text(1,27, expression("LOD"["95%"]*""), cex=1.3, col="grey70")
text(1.1,24.5, expression("C"[q]* " Cut off = 35.25"), cex=1.3, col="grey70")
text(1,9, expression("Experimental"), cex=1.3)
text(1,7, expression("C"[q]* " Cut off = 35.7"), cex=1.3)

par(xpd=NA)
segments(c(p95_log10CN, p95_log10CN, -.28, p95_log10CN), c(lod95cq,lod95cq-12,lod95cq, 6 ), c(p95_log10CN, p95_log10CN,p95_log10CN, p95_log10CN ), c(lod95cq-7, 10,lod95cq, -1.5 ), lty=3, lwd=2, col="grey70")
segments(c(-0.28,.6616,.6616, .6616),c(35.7,35.7,23, 6),c(.6616,.6616,.6616, .6616),c(35.7,28.25,10,-1.5), lwd=2)
par(xpd=F)
dev.off()
setwd(unlist(strsplit(getwd(), "/Figures"))[1])

#####Data oospores extractions

#Load data
setwd(paste(getwd(), "Data", sep="/"))

data_os=read.csv("Oospores_data.csv", header=T)
setwd(unlist(strsplit(getwd(), "/Data"))[1])

#split data into soil and spores
os_soil=subset(data_os, data_os$Sample_type=="Soil")
os_soil[,5]=as.numeric(os_soil[,5])

os_spores=subset(data_os, data_os$Sample_type=="Spores")
os_spores[,5]=as.numeric(os_spores[,5])


####set the lower boundary and higher boundary for the outliers filtering

spocount=unique(os_soil$Total_Spore_number)
bound_s=data.frame(cell_num=double(), median=double(), higher=double(), lower=double(), obs=double())
bound_sp=data.frame(cell_num=double(), median=double(), higher=double(), lower=double(), obs=double())

out_tab_s=data.frame(spore_num=double(), potential_outliers=character())
out_tab_sp=data.frame(spore_num=double(), potential_outliers=character())

####determine the median and the median absolute deviation and determine which
####data points can be considered outliers.

for (i in spocount){

	if(i==spocount[1]){
		l=1
	}else{
		l=l+1
	}

	subs=os_soil[os_soil$Total_Spore_number==i,5]
	subsp=os_spores[os_soil$Total_Spore_number==i,5]

	obs_s=length(na.omit(subs))
	obs_sp=length(na.omit(subsp))
	median_s=median(na.omit(subs))
	median_sp=median(na.omit(subsp))
	lower_s=median_s-(3*mad(na.omit(subs), constant=1))
	lower_sp=median_sp-(3*mad(na.omit(subsp), constant=1))
	higher_s=median_s+(3*mad(na.omit(subs), constant=1))
	higher_sp=median_sp+(3*mad(na.omit(subsp), constant=1))
	
	bound_s[l,c(1:5)]=c(i, median_s, higher_s, lower_s, obs_s)
	bound_sp[l,c(1:5)]=c(i, median_sp, higher_sp,  lower_sp, obs_sp)

	out_s=which(subs<lower_s | subs> higher_s)
	out_sp=which(subsp<lower_sp | subsp> higher_sp)

	if(length(out_s)==0){
		out_tab_s[l,c(1:2)]=c(i, "NA")
	}else{
		outlist=paste(list(out_s[]))
		out_tab_s[l,c(1:2)]=c(i,paste(list(subs[out_s[1:length(out_s)]])))
	}

	if(length(out_sp)==0){
		out_tab_sp[l,c(1:2)]=c(i, "NA")
	}else{
		outlist=paste(list(out_sp[]))
		out_tab_sp[l,c(1:2)]=c(i,paste(list(subsp[out_sp[1:length(out_sp)]])))
	}

} 

##exclude data identified by outlier analysis###
os_soil_cur=os_soil[-c(40,66,71, 106),]
os_spores_cur=os_spores[-c(37,38,39,42,58,59,60,63,68,82,83),]

#remove NAs from the table to allow for mean
os_soil_cur=na.omit(os_soil_cur)
os_spores_cur=na.omit(os_spores_cur)


#calculate the mean of each Cq for individual biological replicate
dp_soil=aggregate(os_soil_cur[,5], by=list(os_soil_cur[,1], os_soil_cur[,3]), FUN=mean)
colnames(dp_soil)=c("spore_num", "bio_rep", "Cq")
dp_soil$Log_spores_per_gram=log(dp_soil[,1]/.25,10)

dp_spores=aggregate(os_spores_cur[,5], by=list(os_spores_cur[,1], os_spores_cur[,3]), FUN=mean)
colnames(dp_spores)=c("spore_num", "bio_rep", "Cq")
dp_spores$Log_spores=log(dp_spores[,1]/.25, 10)

###remove the data points lower than 8.3
dp_soil_final=dp_soil[-c(1,2,3,8,9,10,15,16,21,22),]
dp_spores_final=dp_spores[-c(1,6,7,8,14,15,16),]


####keep the removed points to put into graph as removed data
dp_soil_rm=dp_soil[c(1,2,3,8,9,10,15,16,21,22),]
dp_spores_rm=dp_spores[c(1,6,7,8,14,15,16),]


##estimate the standard curve with a linear model
soil_lm=lm(dp_soil_final[,3]~dp_soil_final[,4])
rsq_soil=round(cor(dp_soil_final[,4], dp_soil_final[,3])^2, 3)
slope_soil=round(soil_lm$coefficients[2],3)
ordori_soil=round(soil_lm$coefficients[1],3)

spores_lm=lm(dp_spores_final[,3]~dp_spores_final[,4])
rsq_spores=round(cor(dp_spores_final[,4], dp_spores_final[,3])^2, 3)
slope_spores=round(spores_lm$coefficients[2],3)
ordori_spores=round(spores_lm$coefficients[1],3)


###create data from the standard curve equations
pts_s=c(1.521138:4.521138)
pts_sp=c(1.380211,1.521138:4.521138)
soilcurve=pts_s*soil_lm$coefficients[2]+soil_lm$coefficients[1]
sporescurve=pts_sp*spores_lm$coefficients[2]+spores_lm$coefficients[1]


#plot of the Cq in function on the log10 spore
setwd(paste(getwd(), "Figures", sep="/"))
png("Figure_1D.png", height=2000, width=2000, res=300)
par(mar=c(5,5,4,1))
plot.new()
plot.window(xlim=c(0,5), ylim=c(10,40))
axis(1, lwd=2, cex.axis=1.5)
axis(2, lwd=2, cex.axis=1.5, las=1)
title(xlab=expression("log"[10]*" Oospore number"), cex.lab=1.5, ylab=expression("C"[q]*" Value"))
points(dp_soil_final[,4], dp_soil_final[,3], pch=16, col=rgb(0,0,0,1), cex=1.5)
points(dp_spores_final[,4], dp_spores_final[,3], pch=16, col=rgb(0,0,0,.25), cex=1.5)

points(dp_spores_rm[,4], dp_spores_rm[,3], col=rgb(1,0,0,.25), pch=17, cex=1.5)
points(dp_soil_rm[,4],dp_soil_rm[,3], col=rgb(1,0,0,1), pch=17, cex=1.5)

lines(pts_s,soilcurve, lwd=2, lty=2, col=rgb(0,0,0,1))
lines(pts_sp,sporescurve, lwd=2, lty=2, col=rgb(0,0,0,.5))

text(4,36, bquote(paste("R"["2"]*" = ",.(rsq_soil))), cex=1.3) 
text(4,39, bquote(paste("C"["q"]* " = ", .(slope_soil), "x + ",.(ordori_soil))),  cex=1.3)

text(4,30, bquote(paste("R"["2"]*" = ",.(rsq_spores))), cex=1.3, col=rgb(0,0,0,.5)) 
text(4,33, bquote(paste("C"["q"]* " = ", .(slope_spores), "x + ",.(ordori_spores))),  cex=1.3, col=rgb(0,0,0,.5))

legend(1.25,17, c("Oospores with soil", "Oospores alone", "removed data"), pch=c(16,16,17), lty=c(2,2,0), lwd=2, col=c(rgb(0,0,0,1), rgb(0,0,0,.5), rgb(1,0,0,1)), bty="n", cex=1.5)
dev.off()

setwd(unlist(strsplit(getwd(), "/Figures"))[1])


###compare the regression lines

#pool data into one set and Identify them with either soil or spores
dp_all=as.data.frame(rbind(cbind(dp_soil_final[,3],dp_soil_final[,4]), cbind(dp_spores_final[,3], dp_spores_final[,4])))
colnames(dp_all)=c("Cq", "Log10_spores")
dp_all$Sample_type=cbind(c(rep("Soil",16), rep("Spores",18)))

#make a new regression analysis with the sample type as an independent variable
sospo_lm=lm(Cq~Log10_spores+Sample_type+Sample_type:Log10_spores, data=dp_all)

summary(sospo_lm)

#Coefficients:
#                               Estimate Std. Error t value Pr(>|t|)    
#(Intercept)                     34.2659     0.7688  44.572  < 2e-16 ***
#Log10_spores                    -2.9521     0.2386 -12.370  2.6e-13 ***
#Sample_typeSpores                2.4478     1.0084   2.427   0.0214 *  
#Log10_spores:Sample_typeSpores  -0.6013     0.3197  -1.881   0.0697 .  
#---
#Signif. codes:  0 �***� 0.001 �**� 0.01 �*� 0.05 �.� 0.1 � � 1
#
#Residual standard error: 1.067 on 30 degrees of freedom
#Multiple R-squared:  0.9368,    Adjusted R-squared:  0.9305 
#F-statistic: 148.3 on 3 and 30 DF,  p-value: < 2.2e-16


########################################
########Data analysis field data########
########################################

setwd(paste(getwd(), "/Data", sep=""))
Data=read.csv("Data_qPCR_Pabi.csv", header=T)
metadata=read.csv("metadata_phytophthora_2019_2021.csv", header=T)
setwd(unlist(strsplit(getwd(),"/Data"))[1])

#create a vector with all the sample names and types
Sample_names=unique(Data$sample)
Sample_types=unique(Data$sample_type)

###Create a results table with column for sample, Cq average, Detection status
###and number of replicates Res_table_dataed

Res_table=data.frame(Sample=character(), Sample_type=character(), Cq_average=double(), 
Detection=integer(), Replicates=double(), Flag=character())

####start at line 1 in the new data frame since S is the first sample type
for (type in Sample_types){

	if(type=="S"){
	line=1
	}else{
	line=line
	}

	for (samp in Sample_names){

#set threshold to desired value
		det_threshold=35.7
#sub is a vector of the 3 replicates Ct values
		sub=subset(Data$Cq_Pabi, Data$sample == samp & Data$sample_type == type)
#calculate the number of replicates
		repnum=length(sub)
#calculate the number of "Undetermined" replicate values for current sample
		undet_num=length(which(sub=="Undetermined"))
#calculate the number of replicate over the threshold value for current sample
		thresh_fail=length(which(sub > det_threshold))
#calculate the number of NA
		na_num=length(which(is.na(sub)))
#if there is more than one "Undetermined" or Ct over threshold the
#average will be NA and there is no detection
			if(undet_num > 0 | thresh_fail > 0 | na_num > 0){
				avg="NA"
				det=0
#in the other cases, the average Ct will be calculated and there is detection
			}else{
				avg=round(mean(as.numeric(sub)), 2)
				det=1
			}
#if the number of replicates is different from 3 --> flag sample
			if(repnum!=3){
				flag="*"
			}else{
				flag=""
			}

#Report the information to the Res_table at the right line
#if the number of replicate is 0 skip this step
			if (repnum == 0){
				next
			}else{
		Res_table[line,]= c(samp, type,avg, det, repnum, flag)
		line=line+1
		}
	}
}

##merge to metadata file
Res_table_data=merge(metadata, Res_table)

###change columns to the right class (numeric, factor...etc)
Res_table_data$Sample=as.numeric(Res_table_data$Sample)
Res_table_data$Date=as.factor(Res_table_data$Date)
Res_table_data$Year=as.factor(Res_table_data$Year)
Res_table_data$Region=as.factor(Res_table_data$Region)
Res_table_data$Species=as.factor(Res_table_data$Species)
Res_table_data$Status=as.factor(Res_table_data$Status)
Res_table_data$Baiting_Phytophthora_isolation=as.factor(Res_table_data$Baiting_Phytophthora_isolation)
Res_table_data$Baiting_P_abietivora_isolation=as.factor(Res_table_data$Baiting_P_abietivora_isolation)
Res_table_data$Root_Phytophthora_isolation=as.factor(Res_table_data$Root_Phytophthora_isolation)
Res_table_data$Root_P_abietivora_isolation=as.factor(Res_table_data$Root_P_abietivora_isolation)
Res_table_data$Sample_type=as.factor(Res_table_data$Sample_type)
Res_table_data$Cq_average=as.numeric(Res_table_data$Cq_average)
Res_table_data$Detection=as.factor(Res_table_data$Detection)
Res_table_data$Replicates =as.numeric(Res_table_data$Replicates)


###write .csv file with results in a new directory and comback to the Data directory
dir.create(paste(getwd(), "Results", sep="/"))
setwd(paste(getwd(), "Results", sep="/"))
write.table(Res_table_data,"Detection_qPCR_Pabi.csv" ,sep=",", row.names=F)
setwd(unlist(strsplit(getwd(), "/Results"))[1])


######Create subset for soil and root samples
Soil=subset(Res_table_data, Res_table_data$Sample_type == "S")
row.names(Soil)=c(1:nrow(Soil))
Roots=subset(Res_table_data, Res_table_data$Sample_type == "R")
row.names(Roots)=c(1:nrow(Roots))

################Positive/Negative Soil samples data tables##########################
soil_spec=table(Soil$Species, Soil$Detection)
#                      0   1
#      Abies balsamea 308  60
#      Abies fraseri  112   8

table_species=cbind(soil_spec[,2], soil_spec[,1])
t_soil_spec=prop.test(table_species)

#2-sample test for equality of proportions with continuity correction

#data:  table_species
#X-squared = 6.2281, df = 1, p-value = 0.01257
#alternative hypothesis: two.sided
#95 percent confidence interval:
# 0.03240197 0.16035166
#sample estimates:
#    prop 1     prop 2 
#0.16304348 0.06666667        

soil_stat=table(Soil$Status, Soil$Detection)                    
#                       0   1
#  Healthy_forest     151   9
#  Healthy_plantation 147  17
#  PRR_symptomatic    122  42

table_status=cbind(soil_stat[,2], soil_stat[,1])
t_soil_stat=pairwise.prop.test(table_status, p.adjust.method="fdr")

#Pairwise comparisons using Pairwise comparison of proportions 
#data:  table_status 
#                   Healthy_forest Healthy_plantation
#Healthy_plantation 0.17197        -                 
#PRR_symptomatic    5.1e-06        0.00084   


###barplot proportion of positive in sample health status
setwd(paste(getwd(), "Figures", sep="/"))

png("Figure_3A.png", height=2000, width=2200, res=300)
par(mar=c(7,5,4,2))
barplot(t(prop.table(soil_stat,1)), lwd=2, axes=F, cex.names=1.2, border=NA, col=c(rgb(0,0,0,1), rgb(0,0,0,.35)), names.arg=c("Healthy forest", "Healthy plantation", "PRR symptomatic"))
axis(2,las=2, lwd=2, cex.axis=1.5)
title(xlab="Sample type", cex.lab=1.5)
title(ylab="Proportion", cex.lab=1.5, line=3.5)
par(xpd=NA)
legend(1.2,-.2, c("Negative", "Positive"), fill=c(rgb(0,0,0,1), rgb(0,0,0,.35)), horiz=T, border=NA)
text(c(0.7,1.9,3.1), 1.05, c("a","a","b"), cex=1.8)
par(xpd=F)
dev.off()

setwd(unlist(strsplit(getwd(), "/Figures"))[1])


soil_reg=table(Soil$Region, Soil$Detection) 
#                        0   1
#  Chaudiere_Appalache 206  42
#  Estrie              214  26

table_reg=cbind(soil_reg[,2], soil_reg[,1])
t_soil_reg=prop.test(table_reg)

#2-sample test for equality of proportions with continuity correction
#
#data:  table_reg
#X-squared = 3.2952, df = 1, p-value = 0.06948
#alternative hypothesis: two.sided
#95 percent confidence interval:
# -0.004111921  0.126154932
#sample estimates:
#   prop 1    prop 2 
#0.1693548 0.1083333 


soil_year=table(Soil$Year, Soil$Detection)
#         0   1
#  2019 212  28
#  2020 101  27
#  2021 107  13

table_year=cbind(soil_year[,2], soil_year[,1])
t_soil_year=pairwise.prop.test(table_year, p.adjust.method="fdr")

#Pairwise comparisons using Pairwise comparison of proportions 

#data:  table_year 

#     2019  2020 
#2020 0.065 -    
#2021 0.953 0.065
#
#P value adjustment method: fdr 

################Positive/Negative root samples data tables##########################

root_spec=table(Roots$Species, Roots$Detection)
#                     0   1
#     Abies balsamea 103  19
#     Abies fraseri   37   9

table_species_r=cbind(root_spec[,2], root_spec[,1])
prop.test(table_species_r)

#2-sample test for equality of proportions with continuity correction
#
#data:  table_species_r
#X-squared = 0.14968, df = 1, p-value = 0.6988
#alternative hypothesis: two.sided
#95 percent confidence interval:
# -0.1863443  0.1065154
#sample estimates:
#   prop 1    prop 2 
#0.1557377 0.1956522 


root_reg=table(Roots$Region, Roots$Detection) 
#                       0  1
#  Chaudiere_Appalache 73 15
#  Estrie              67 13

table_reg_r=cbind(root_reg[,2], root_reg[,1])
prop.test(table_reg_r)

#2-sample test for equality of proportions with continuity correction
#
#data:  table_reg_r
#X-squared = 1.8138e-30, df = 1, p-value = 1
#alternative hypothesis: two.sided
#95 percent confidence interval:
# -0.1127275  0.1286366
#sample estimates:
#   prop 1    prop 2 
#0.1704545 0.1625000 
 

root_year=table(Roots$Year, Roots$Detection)
#        0  1
#  2019 67  13
#  2020 38 10
#  2021 35  5

table_year=cbind(root_year[,2], root_year[,1])
pairwise.prop.test(table_year, p.adjust.method="fdr")

#Pairwise comparisons using Pairwise comparison of proportions 
#
#data:  table_year 
#
#     2019 2020
#2020 0.79 -   
#2021 0.79 0.79


##########Map of regions with different samplings##########

setwd(paste(getwd(), "/Data", sep=""))

sampling_area=st_read("regio_s.shp")

setwd(unlist(strsplit(getwd(),"/Data"))[1])

Soil$Coordinates=paste(Soil$Latitude, Soil$Longitude, sep=",")
sites=unique(Soil$Coordinates)

#map quebec with the study regions in white
setwd(paste(getwd(), "Figures", sep="/"))
png("Figure_2A.png", height=2200, width=2200, res=300)
par(mar=c(5,7,4,4))
plot.new()
plot.window(xlim=c(-80,-57), ylim=c(44,63))
plot(sampling_area, lwd=2, col=c(rep("grey70", 12), rgb(0/255,0/255,0/255,0), rep("grey70", 2), rgb(0/255,0/255,0/255,0), rep("grey70", 5)), add=T)
axis(1, at=seq(-80,-57, 5.75), cex.axis=1.7, lwd=3)
axis(2, at=seq(44,63, 4.75), lwd=3, cex.axis=1.7, las=1)
title(xlab="longitude", cex.lab=2)
title(ylab="latitude", cex.lab=2, line=5)
dev.off()


###PRR symtomatic trees####

png("Figure_2B.png", height=2200, width=2200, res=300)
par(mar=c(5,7,4,4))
plot.new()
plot.window(xlim=c(-73.05,-69.75), ylim=c(45,47.25))
plot(sampling_area[,7], lwd=2.5, col=c(rep("grey70", 12), rgb(0/255,0/255,0/255,0), rep("grey70", 2), rgb(0/255,0/255,0/255,0), rep("grey70", 5)), add=T)
axis(1, at=seq(-73.75,-69.5, .75), cex.axis=1.7, lwd=3)
axis(2, at=seq(45.05,48.05, .75), lwd=3, cex.axis=1.7, las=1)
title(xlab="longitude", main="PRR symptomatic trees", cex.lab=2, cex.main=2)
title(ylab="latitude", cex.lab=2, line=5)


for (i in 1:length(sites)){

	sub=subset(Soil, Soil$Coordinates == sites[i] & Soil$Status == "PRR_symptomatic")
	lat=strsplit(sites, ",")[[i]][1]
	long=strsplit(sites, ",")[[i]][2]
	pt_col=length(which(sub$Detection ==1)== TRUE)/nrow(sub)
	if(pt_col==0){
		points(long, lat, col="black", pch=13, cex=1.5)
	}else{
		points(long, lat, col="black", pch=pt_col*8, cex=1.5)
}
}

legend("topleft", c("0",".125", ".250", ".375", ".500", ".625", ".750", ".875", "1.00"), cex=1.25, pt.cex=1.5, bg="white", 
col="black" ,pch=c(13,1,2,3,4,5,6,7,8), box.lwd=2, title="Proportion of positive detection", ncol=3)

dev.off()

######inset beauce

png("Figure_2B_inset.png", height=2200, width=2200, res=300)
par(mar=c(3,4.5,2,2))
plot.new()
plot.window(xlim=c(-71,-70.75), ylim=c(45.8,46.2))
plot(sampling_area[,7], lwd=2.5, col=c(rep("grey90", 12), rgb(140/255,193/255,64/255,0), rep("grey90", 2), rgb(140/255,193/255,64/255,0), rep("grey90", 5)), add=T)
axis(1, at=seq(-71,-70.75, .125), cex.axis=2, lwd=3)
axis(2, at=seq(45.8,46.2, .1), lwd=3, cex.axis=2, las=1)


for (i in 1:length(sites)){

	sub=subset(Soil, Soil$Coordinates == sites[i] & Soil$Status == "PRR_symptomatic")
	lat=strsplit(sites, ",")[[i]][1]
	long=strsplit(sites, ",")[[i]][2]
	pt_col=length(which(sub$Detection ==1)== TRUE)/nrow(sub)
	if(pt_col==0){
		points(long, lat, col="black", pch=13, cex=3)
	}else{
		points(long, lat, col="black", pch=pt_col*8, cex=3)
}
}

dev.off()
####Healthy plantation trees####
png("Figure_2C.png", height=2200, width=2200, res=300)
par(mar=c(5,7,4,4))
plot.new()
plot.window(xlim=c(-73.05,-69.75), ylim=c(45,47.25))
plot(sampling_area[,7], lwd=2.5, col=c(rep("grey70", 12), rgb(0/255,0/255,0/255,0), rep("grey70", 2), rgb(0/255,0/255,0/255,0), rep("grey70", 5)), add=T)
axis(1, at=seq(-73.75,-69.5, .75), cex.axis=1.7, lwd=3)
axis(2, at=seq(45.05,48.05, .75), lwd=3, cex.axis=1.7, las=1)
title(xlab="longitude", main="Healthy trees in plantation", cex.lab=2, cex.main=2)
title(ylab="latitude", cex.lab=2, line=5)


for (i in 1:length(sites)){

	sub=subset(Soil, Soil$Coordinates == sites[i] & Soil$Status == "Healthy_plantation")
	lat=strsplit(sites, ",")[[i]][1]
	long=strsplit(sites, ",")[[i]][2]
	pt_col=length(which(sub$Detection ==1)== TRUE)/nrow(sub)
	if(pt_col==0){
		points(long, lat, col="black", pch=13, cex=1.5)
	}else{
		points(long, lat, col="black", pch=pt_col*8, cex=1.5)
}
}

legend("topleft", c("0",".125", ".250", ".375", ".500", ".625", ".750", ".875", "1.00"), cex=1.25, pt.cex=1.5, bg="white", 
col="black" ,pch=c(13,1,2,3,4,5,6,7,8), box.lwd=2, title="Proportion of positive detection", ncol=3)

dev.off()

###Inset Beauce
png("Figure_2C_inset.png", height=2200, width=2200, res=300)
par(mar=c(3,4.5,2,2))
plot.new()
plot.window(xlim=c(-71,-70.75), ylim=c(45.8,46.2))
plot(sampling_area[,7], lwd=2.5, col=c(rep("grey90", 12), rgb(140/255,193/255,64/255,0), rep("grey90", 2), rgb(140/255,193/255,64/255,0), rep("grey90", 5)), add=T)
axis(1, at=seq(-71,-70.75, .125), cex.axis=2, lwd=3)
axis(2, at=seq(45.8,46.2, .1), lwd=3, cex.axis=2, las=1)

for (i in 1:length(sites)){

	sub=subset(Soil, Soil$Coordinates == sites[i] & Soil$Status == "Healthy_plantation")
	lat=strsplit(sites, ",")[[i]][1]
	long=strsplit(sites, ",")[[i]][2]
	pt_col=length(which(sub$Detection ==1)== TRUE)/nrow(sub)
	if(pt_col==0){
		points(long, lat, col="black", pch=13, cex=3)
	}else{
		points(long, lat, col="black", pch=pt_col*8, cex=3)
}
}

dev.off()

####Healthy forest trees####
png("Figure_2D.png", height=2200, width=2200, res=300)
par(mar=c(5,7,4,4))
plot.new()
plot.window(xlim=c(-73.05,-69.75), ylim=c(45,47.25))
plot(sampling_area[,7], lwd=2.5, col=c(rep("grey70", 12), rgb(0/255,0/255,0/255,0), rep("grey70", 2), rgb(0/255,0/255,0/255,0), rep("grey70", 5)), add=T)
axis(1, at=seq(-73.75,-69.5, .75), cex.axis=1.7, lwd=3)
axis(2, at=seq(45.05,48.05, .75), lwd=3, cex.axis=1.7, las=1)
title(xlab="longitude", main="Healthy trees in natural Forest", cex.lab=2, cex.main=2)
title(ylab="latitude", cex.lab=2, line=5)


for (i in 1:length(sites)){

	sub=subset(Soil, Soil$Coordinates == sites[i] & Soil$Status == "Healthy_forest")
	lat=strsplit(sites, ",")[[i]][1]
	long=strsplit(sites, ",")[[i]][2]
	pt_col=length(which(sub$Detection ==1)== TRUE)/nrow(sub)
	if(pt_col==0){
		points(long, lat, col="black", pch=13, cex=1.5)
	}else{
		points(long, lat, col="black", pch=pt_col*8, cex=1.5)
}
}

legend("topleft", c("0",".125", ".250", ".375", ".500", ".625", ".750", ".875", "1.00"), cex=1.25, pt.cex=1.5, bg="white", 
col="black" ,pch=c(13,1,2,3,4,5,6,7,8), box.lwd=2, title="Proportion of positive detection", ncol=3)

dev.off()


###Inset Beauce
png("Figure_2D_inset.png", height=2200, width=2200, res=300)
par(mar=c(3,4.5,2,2))
plot.new()
plot.window(xlim=c(-71,-70.75), ylim=c(45.8,46.2))
plot(sampling_area[,7], lwd=2.5, col=c(rep("grey90", 12), rgb(140/255,193/255,64/255,0), rep("grey90", 2), rgb(140/255,193/255,64/255,0), rep("grey90", 5)), add=T)
axis(1, at=seq(-71,-70.75, .125), cex.axis=2, lwd=3)
axis(2, at=seq(45.8,46.2, .1), lwd=3, cex.axis=2, las=1)

for (i in 1:length(sites)){

	sub=subset(Soil, Soil$Coordinates == sites[i] & Soil$Status == "Healthy_forest")
	lat=strsplit(sites, ",")[[i]][1]
	long=strsplit(sites, ",")[[i]][2]
	pt_col=length(which(sub$Detection ==1)== TRUE)/nrow(sub)
	if(pt_col==0){
		points(long, lat, col="black", pch=13, cex=3)
	}else{
		points(long, lat, col="black", pch=pt_col*8, cex=3)
}
}

dev.off()

setwd(unlist(strsplit(getwd(), "/Figures"))[1])

####glm on positive detection

glm_detection=glm(Detection~Status+Species+Year+Region, data=Soil, family=binomial(logit))

#glm(formula = Detection ~ Status + Species + Year + Region, family = binomial(logit), 
#    data = Soil)

#Deviance Residuals: 
#    Min       1Q   Median       3Q      Max  
#-1.0237  -0.6334  -0.4352  -0.2836   2.4548  
#
#Coefficients:
#                           Estimate Std. Error z value Pr(>|z|)    
#(Intercept)              -2.963e+00  4.281e-01  -6.921 4.48e-12 ***
#StatusHealthy_plantation  6.535e-01  4.320e-01   1.513   0.1303    
#StatusPRR_symptomatic     1.785e+00  3.917e-01   4.557 5.19e-06 ***
#SpeciesAbies fraseri     -1.042e+00  4.178e-01  -2.494   0.0126 *  
#Year2020                  8.047e-01  3.684e-01   2.185   0.0289 *  
#Year2021                  1.956e-15  4.315e-01   0.000   1.0000    
#RegionEstrie              1.575e-01  4.287e-01   0.367   0.7134

##predict the model for the status in Af and Ab while keeping the other variable fixed
stat_ab=data.frame(Status=c("Healthy_forest", "Healthy_plantation", "PRR_symptomatic"), Species=rep("Abies balsamea",3), Year=rep("2019",3), Region=rep("Chaudiere_Appalache",3))
stat_af=data.frame(Status=c("Healthy_forest", "Healthy_plantation", "PRR_symptomatic"), Species=rep("Abies fraseri",3), Year=rep("2019",3), Region=rep("Chaudiere_Appalache",3))


###get the OR and 95% CI for the different variables
exp(cbind(OR=coef(glm_detection), confint.default(glm_detection)))

#get the 95% interval for the status (both in Ab and Af)

SEab=predict(glm_detection, newdata=stat_ab, type="response", se=T)$se.fit
ULab=predict(glm_detection, newdata=stat_ab, type="response")+(2*SEab)
LLab=predict(glm_detection, newdata=stat_ab, type="response")-(2*SEab)
#set the negative values to 0 for the interval
LLab[which(LLab<0)]=0

SEaf=predict(glm_detection, newdata=stat_af, type="response", se=T)$se.fit
ULaf=predict(glm_detection, newdata=stat_af, type="response")+(2*SEaf)
LLaf=predict(glm_detection, newdata=stat_af, type="response")-(2*SEaf)
LLaf[which(LLaf<0)]=0

###figure 3B glm
setwd(paste(getwd(), "Figures", sep="/"))

png("Figure_3B.png", height=2000, width=2000, res=300)
par(mar=c(4,5,4,4))
plot(predict(glm_detection, newdata=stat_ab, type="response"), type="l", col="grey40", lwd=2,
xlim=c(1,3), ylim=c(0,.5), axes=F, xlab="Status", ylab="probability of positive detection", cex.lab=1.5)
axis(1, label=c("Healthy_forest", "Healthy_plantation", "PRR_symptomatic"), at=c(1,2,3), lwd=2, cex.axis=1.2)
axis(2, at=seq(0,0.5, .1), lwd=2, cex.axis=1.2, las=2, line=-.5)
lines(predict(glm_detection, newdata=stat_af, type="response"), type="l", col="grey70", lwd=2)
legend("top",c(expression(italic("Abies balsamea")), expression(italic("Abies fraseri")), "95% CI"), lty=c(1,1,5), lwd=2, col=c("grey40", "grey70", "black"), cex=1.2, bty="n")
lines(c(1,2,3),LLab, lty=5, lwd=2, col="grey40")
lines(c(1,2,3),ULab, lty=5, lwd=2, col="grey40")
lines(c(1,2,3),LLaf, lty=5, lwd=2, col="grey70")
lines(c(1,2,3),ULaf, lty=5, lwd=2, col="grey70")
dev.off()

setwd(unlist(strsplit(getwd(), "/Figures"))[1])

####Cq Data analysis Soil vs Root
###Change the NA values to 40 so the non detection data can be plotted
Soil$Cq_average[is.na(Soil$Cq_average)]=40
Roots$Cq_average[is.na(Roots$Cq_average)]=40

#remove sample from non PRR trees
num_roots_samp=Roots$Sample
num_roots_samp=num_roots_samp[-c(93:98)]


####Select the data for which the we have both the Cq data for roots and Soil 
rootsamp=Roots$Cq_average[which(Roots$Sample%in%num_roots_samp)]
soilsamp=Soil$Cq_average[which(Soil$Sample%in%num_roots_samp)]

#### there are samples (352 and 360) without root Cq data####
missdata=Soil$Cq_average[c(352,360)]

###
det_root_40_soil=which(soilsamp == 40 & rootsamp < 40)
det_soil_40_root=which(soilsamp < 40 & rootsamp == 40)
soil_40_root_40=which(soilsamp == 40 & rootsamp == 40)


###plot the individual datapoints in figure 4A
setwd(paste(getwd(), "Figures", sep="/"))
png("Figure_4A.png", height=2000, width=2000, res=300)
par(xpd=NA, mar=c(5,5,4,4))
plot(soilsamp, rootsamp, axes=F, ylim=c(15,40), xlim=c(15,40),
pch=16, col=rgb(0,0,0,0.25), ylab=expression("C"[q]*" Roots"), xlab=expression("C"[q]*" Soil"), cex.lab=1.5, cex=1.5)
axis(1, lwd=2, cex.axis=1.5)
axis(2, lwd=2, cex.axis=1.5, las=2)
segments(c(39,38.5,38.5),c(23.5,35.5,23.5),c(39,39.5,39.5),c(35.5,35.5,23.5), lwd=2)
segments(c(26,26,35),c(39,39.5,39.5),c(35,26,35),c(39,38.5,38.5), lwd=2)
text(c(30.5,37),c(38, 29), c(paste("n=", length(det_soil_40_root)+2,sep=""), paste("n=", length(det_root_40_soil), sep="")), cex=1.5)
points(40,40, cex=5, lwd=2)
points(missdata,c(41,41), pch=17, cex=1.5, col=rgb(0,0,0,0.25))
text(40,38, paste("n=", length(soil_40_root_40), sep=""), cex=1.5)
legend("bottom", c("data", "missing data for roots"), pch=c(16,17), cex=1.5, col=rgb(0,0,0,.25), bty="n")
par(xpd=FALSE)
dev.off()
setwd(unlist(strsplit(getwd(), "/Figures"))[1])


###test difference in detection proportion
detect_sr=matrix(nrow=2,ncol=2,dimnames=list(c("Soil", "Roots"),c("Positive", "Total")),
data=c(length(which(soilsamp<40))+2,length(which(rootsamp<40)),110+12,110+24))

prop.test(detect_sr)

#2-sample test for equality of proportions with continuity correction

#data:  detect_sr
#X-squared = 2.8748, df = 1, p-value = 0.08998
#alternative hypothesis: two.sided
#95 percent confidence interval:
# -0.01149185  0.17800796
#sample estimates:
#   prop 1    prop 2 
#0.2560976 0.1728395


###Boxplot for Cq values between Soil and Roots samples

ct_PRRs=Res_table_data$Cq_average[which(Res_table_data$Status == "PRR_symptomatic" & Res_table_data$Sample_type == "S")]
ct_PRRr=Res_table_data$Cq_average[which(Res_table_data$Status == "PRR_symptomatic" & Res_table_data$Sample_type == "R")]

setwd(paste(getwd(), "Figures", sep="/"))
png("Figure_4B.png", height=2000, width=2000, res=300)
par(mar=c(4,5,4,4), mfrow=c(1,1))
boxplot(ct_PRRs, ct_PRRr, ylim=c(15,40), axes=F, lty=1, lwd=2)
axis(1, at=c(1,2), c("Soil Samples", "Unidentified Root Fragments"), lwd=2, cex.axis=1.2)
axis(2, at=seq(15, 40,5), lwd=2, cex.axis=1.5)
title(xlab="Sample types", ylab=expression("C"[q]*" Value"), cex.lab=1.5)
text(c(1,2),c(36.5,36.5), c(paste(length(which(ct_PRRs != "NA"))), paste(length(which(ct_PRRr != "NA")))), cex=2)
segments(c(.6,1.6),c(mean(ct_PRRs, na.rm=T), mean(ct_PRRr, na.rm=T)),c(1.4,2.4),c(mean(ct_PRRs, na.rm=T), mean(ct_PRRr, na.rm=T)), lwd=3, lty=3, col=c("black", "white"))
legend("bottom","mean value", lty=3,lwd=3, cex=2, bty="n" )
text(c(1,2),c(39,39),"a", cex=2)
dev.off()
setwd(unlist(strsplit(getwd(), "/Figures"))[1])

####Test the difference in Cq average between roots and soil samples

t.test(ct_PRRs, ct_PRRr)

#Welch Two Sample t-test
#
#data:  ct_PRRs and ct_PRRr
#t = 1.4911, df = 50.197, p-value = 0.1422
#alternative hypothesis: true difference in means is not equal to 0
#95 percent confidence interval:
# -0.3946011  2.6693631
#sample estimates:
#mean of x mean of y 
# 31.22595  30.08857 
```
